# A sectioning and database enrichment approach for improved peptide spectrum matching in large, genome-guided protein sequence databases

**DOI:** 10.1101/843078

**Authors:** Praveen Kumar, James E. Johnson, Caleb Easterly, Subina Mehta, Ray Sajulga, Brook Nunn, Pratik D. Jagtap, Timothy J. Griffin

## Abstract

Multi-omics approaches focused on mass-spectrometry (MS)-based data, such as metaproteomics, utilize genomic and/or transcriptomic sequencing data to generate a comprehensive protein sequence database. These databases can be very large, containing millions of sequences, which reduces the sensitivity of matching tandem mass spectrometry (MS/MS) data to sequences to generate peptide spectrum matches (PSMs). Here, we describe a sectioning method for generating an enriched database for those protein sequences that are most likely present in the sample. Our evaluation demonstrates how this method helps to increase the sensitivity of PSMs while maintaining acceptable false discovery rate statistics. We demonstrate increased true positive PSM identifications using the sectioning method when compared to the traditional large database searching method, whereas it helped in reducing the false PSM identifications when compared to a previously described two-step method for reducing database size. The sectioning method for large sequence databases enables generation of an enriched protein sequence database and promotes increased sensitivity in identifying PSMs, while maintaining acceptable and manageable FDR. Furthermore, implementation in the Galaxy platform provides access to a usable and automated workflow for carrying out the method. Our results show the utility of this methodology for a wide-range of applications where genome-guided, large sequence databases are required for MS-based proteomics data analysis.

## Introduction

Liquid chromatography tandem mass spectrometry (LC-MS/MS) is a high-throughput technique used to identify proteins present in complex biological samples. Experimental MS/MS spectra contain mass-to-charge information on amino acid sequence fragments, which are matched to protein sequences within a database to generate peptide spectrum matches (PSMs). The best PSM assignments are selected based on assigned scores which indicate confidence of the MS/MS spectrum matched to a corresponding peptide sequence within the database^1–4^. Scoring of PSM matches depends on factors like the fragmentation quality and signal-to-noise ratio, as well as inclusion of post-translational modifications, and composition of the sequences contained in the database, including the number of sequences^4^. Peptide sequences from PSMs are ultimately assigned to proteins or protein groups by using protein inference methods.

For conventional single-organism, mass-spectrometry (MS)-based proteomic experiments, a reference database containing all known and validated proteins from the organism are used for matching to MS/MS spectra. However, recently newer approaches, which utilize genomics and /or transcriptomics information, have emerged that utilize customized protein sequence databases^5, 6^. One such approach is metaproteomics^6–8^. Metaproteomics employs LC-MS/MS to generate data for identification of proteins expressed by a microbial community. It offers extensive and conclusive inferences about the taxonomic composition and functional impacts of the microbial community and its surroundings (e.g. host organism, environmental ecosystem)^9^. Metaproteomics studies are actively applied in studying the microbiome of environmental ecosystems^9–12^, and also for investigating the microbiome contained in the gut^13, 14^, oral cavity^15^, lavage^16^, and other sites from humans and animal-models^13–19^.

For metaproteomics studies of complex communities of microorganisms, wherein a reference protein sequence database is either not available or is incomplete, data-processing methods need to be employed to generate a customized database. For example, metagenomic and/or metatranscriptomic sequencing data from same or related samples can be used to generate a customized database of proteins that may be expressed by the community^20–24^. For such an approach, methods have been described for protein sequence database generation such as SixGill^20^, MOCAT^21, 22^ and Graph2Pro^23, 24^. Additionally, if the taxonomic composition of a sample is known, a composite database can be created which includes all of the reference protein sequences for the organisms thought to be present in the sample^25, 26^.

Regardless of the method used for sequence database generation in metaproteomics, it generally generates large databases containing orders of magnitude more protein sequences than those used for more conventional, single-organism studies. Although these proteins are meant to enable more comprehensive identification of proteins, such large databases increase potential for PSMs that are “*close-but-not-perfect,*” thus increasing false positive identifications^27, 28^. A target-decoy database approach^29, 30^ is usually used to estimate the false discovery rate (FDR) and control the false identifications. Unfortunately, in controlling for false positives with increased database size more stringent score thresholds are needed, which decreases the number of qualifying true PSM identifications, effectively decreasing sensitivity of identifying peptides truly contained in the sample^28^. For example, Kumar et al. have demonstrated, using a *Mycobacterium tuberculosis* database, that increasing the protein database size can result in the decrease of true PSM identifications^28^. Thus, it is recommended to create databases that balance the composition of proteins potentially present in the sample with database size so that it still maximizes true positive PSMs.

Researchers have suggested some approaches in order to address the challenges in using large sequence databases for metaproteomics studies. These data processing methods generally seek to decrease the size of the database used when matching MS/MS to sequences to increase the sensitivity for true positive PSMs, and include described approaches such as the multi-stage method and two-step method^31–33^. Jagtap et al. proposed a two-step database searching method to address the issue of large databases^33^ demonstrating that the method could aid in increasing the number of true positive PSMs. In this method, MS/MS are first searched against the large database, and PSMs are accepted using very low stringency scoring in order to infer proteins possibly present, and create a smaller, enriched database of proteins most likely present in the sample. This enriched database is then matched to MS/MS data in a second step of sequence database searching. The two-step method has gained acceptance and has been used widely in metaproteomics studies^15, 24, 34^.

Despite its value, concerns about the potential for increased false positives and the need for validation, acknowledged by the original authors^33^ and others^35, 36^ have been raised when using this traditional two-step method. Using benchmark datasets and an entrapment database^37^, we have observed that the reduction of the database size increases the number of total PSMs. However, concern exists that the method biases the composition of the enriched database leading to increased potential for false-positives^35^. Although suggested by others^37^, an in-depth evaluation of this possible shortcoming of the two-step method has not been carried out to-date.

In this work, we sought to more deeply evaluate and modify the traditional two-step method, and develop a method that overcomes its limitations, specifically the inherent potential for increased false positive PSMs. As such, we have developed a database sectioning method [**Figure 1**], wherein a large database is randomly divided into “*n*” number of smaller subset databases. Each sectioned subset database is searched against the complete LC-MS/MS dataset using the target-decoy method^29^. Matches from each of the searches are used to create a smaller database, enriched for protein sequences most likely present in the sample, while still retaining an adequate number of random noise sequences not present in the sample to control the false positive rate. This enriched database is matched to the entire MS/MS dataset in a second step, using the target-decoy method^29^. We have evaluated the sectioning method using two benchmarking datasets and entrapment databases. Our results show that the sectioning method finds a middle ground between the two commonly used methods in metaproteomics, traditional large database search and traditional two-step database search, balancing sensitivity for true PSM identifications while controlling the rate of false-positive PSMs. We also demonstrate the improvements in results when using the sectioning method on large, previously characterized metaproteomic datasets. The method has been developed in the Galaxy for proteomics (Galaxy-P) platform, which will allow for accessibility and flexibility in its use for a wide-variety of applications where large sequence databases are necessary.

**Figure 1:**
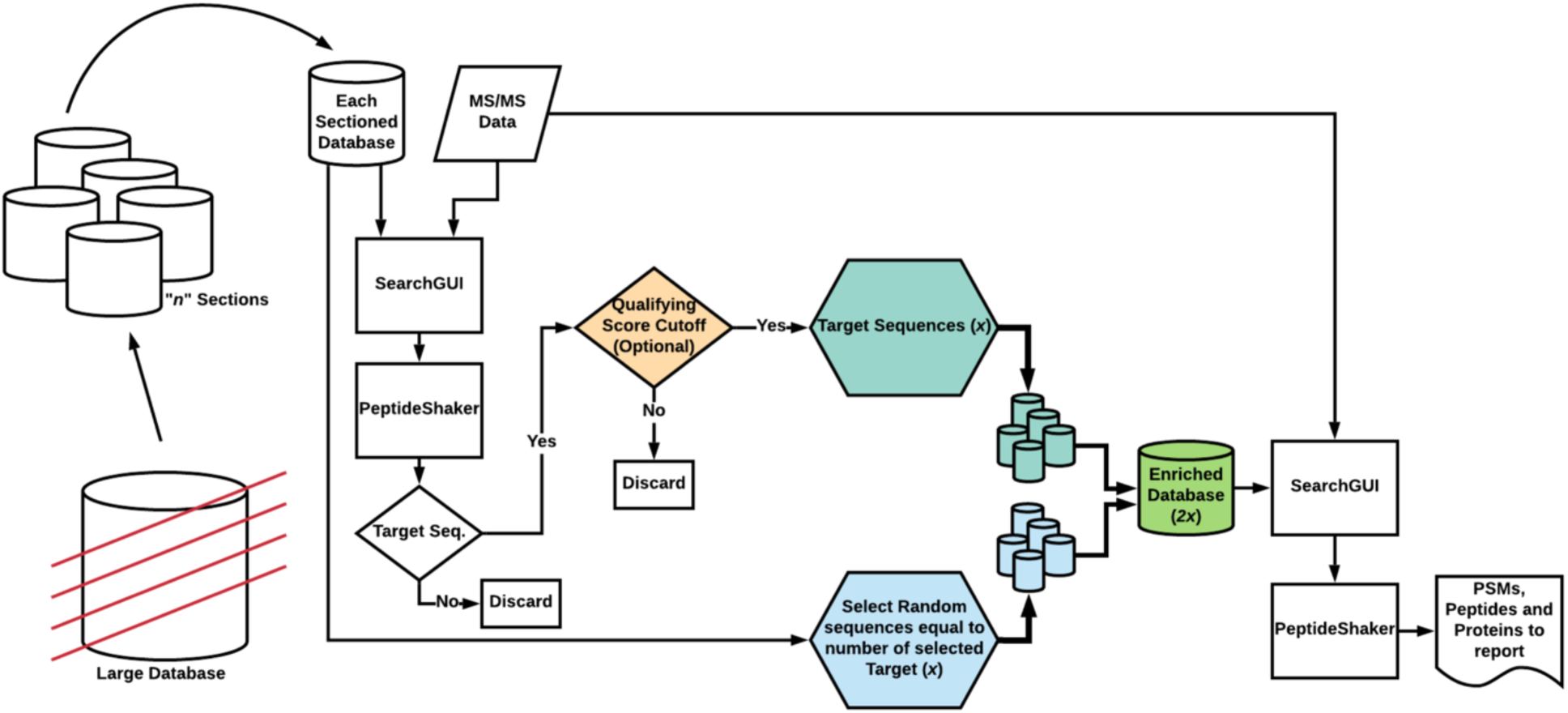
Overview of sectioning method

## Results

To evaluate the effectiveness of our sectioning method, we compared it to the PSM output from two other methods: traditional sequence database searching against the complete protein sequence database (Traditional large database search) and the two-step method for metaproteomics that we originally described (Traditional two-step)^33^ [**Figure 2**]. These three methods all used an entrapment database to help estimate the rate of false positive PSMs compared to true-positive PSMs (see Methods section). In this evaluation, a standard PSM report used as a baseline result where the MS/MS dataset was searched against the database containing sequences from organisms known to be present in the sample. These baseline results were used as a “gold standard” to compare to results when using a metaproteomic database containing sequences from many organisms, some of which may not be present in the sample. Organism-specific proteome(s) sequence databases are recommended for optimal results in MS-based proteomic studies^28^.

**Figure 2:**
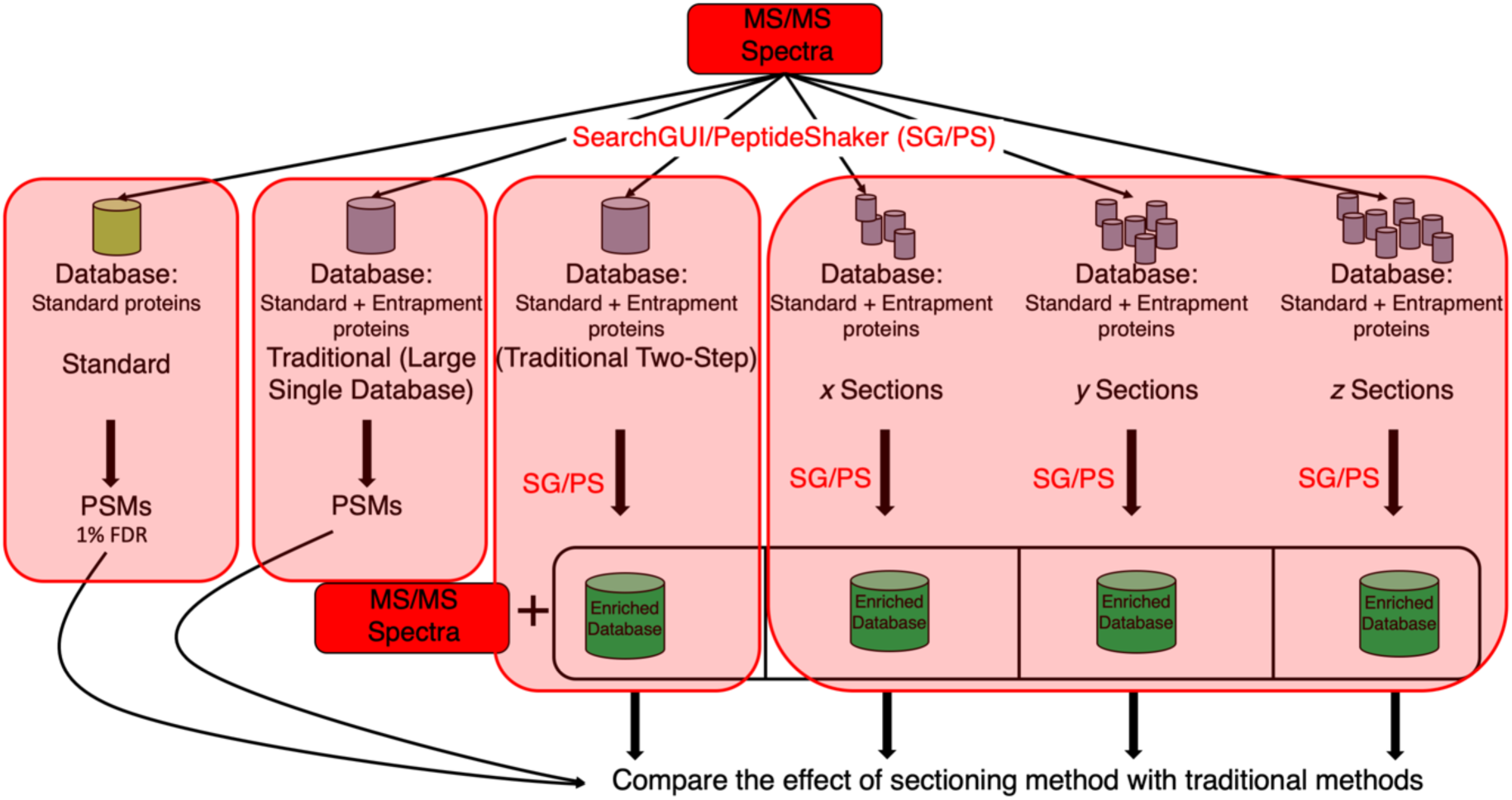
Overview of evaluation method. For each evaluation dataset, the MS/MS spectra were first matched to peptides within a database containing only sequences from organisms known to be present in the sample. Then, using a larger entrapment database, the MS/MS were matched to peptides using the traditional large sequence database searching method, the traditional two-step method, or the sectioning method, using differing numbers of database sections of equal size.

### Pyrococcus furiosus dataset

As a starting point for evaluating our sectioning method, a MS/MS dataset from *Pyrococcus furiosus* (*Pfu*) was matched against a reference *Pfu* proteome database downloaded from Uniprot and contaminant sequences. At 1% global FDR estimated by standard target-decoy methods, this analysis yielded 10,770 PSMs using the standard database. Traditional large database search, traditional two-step method, and the sectioning methods all used entrapment database that contained *Pfu* proteome sequences, common contaminants, and human proteome sequences. *Pfu* has been established as an optimal organism for evaluating PSM results from software algorithms^37, 38^, as its proteome sequence is almost completely orthogonal to higher organisms (e.g. human). Therefore, when using human protein sequences as the entrapment database, true-positive PSMs (PSMs to *Pfu* sequences) can easily be distinguished from false positive PSMs (PSMs to human sequences). This ground truth data can be used to estimate a more accurate FDR compared to the global FDR estimated by target-decoy methods. Here, the entrapment database was comprised of 95601 protein sequences from human primarily, along with the common contaminant protein sequences commonly found in MS-based proteomics experiments. We considered any PSMs identified from *Pfu* or contaminant proteins as true PSMs whereas those identified from human proteins as false PSMs, as per the design of the experiment^37, 38^.

By using this entrapment database for traditional large database search, a decline of 502 PSMs in the sensitivity was observed when the results were compared with the standard search results against the *Pfu* database only (no human entrapment sequence) [**Figure 3a**]. On the other hand, the traditional two-step method identified 10,621 true PSMs to *Pfu* sequences, which recovered the sensitivity and was only 149 PSMs less than the standard search results against the *Pfu* database only [**Figure 3a**].

**Figure 3:**
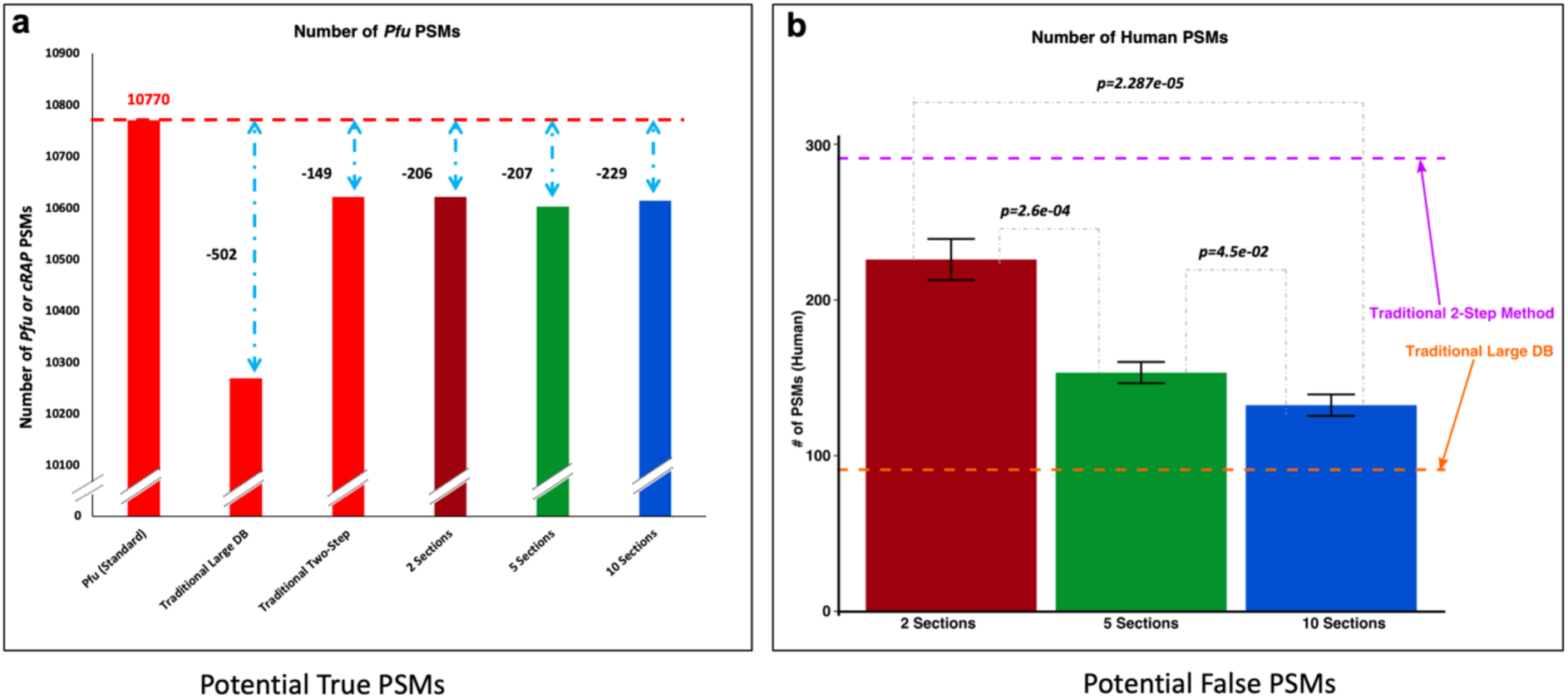
Observations using *Pfu* data. We considered PSMs that matched to a *Pfu* or cRAP protein as true, whereas those matched exclusively to entrapment Human proteins were considered as false PSMs. (a) 10770 PSMs were identified when LC-MS/MS data was matched against *Pfu + cRAP* database. We observed a drop in true PSMs when LC-MS/MS data was matched against an entrapment database (*Pfu* + cRAP + Human protein sequences) using the Traditional Large database search. The Traditional Two-Step method recovered most of the lost PSMs, as well as the sectioning method. (b) The Traditional Large database search controlled the observed false identifications (PSMs matching to Human protein) at a rate of 1%, consistent with the 1% global FDR threshold (orange dashed line). On the contrary, The Traditional Two-Step method inflated the number of false PSMs identified (purple dashed line). Meanwhile, using the sectioning method, we observed decrease in the false PSM identifications when compared with the traditional two-step method (colored bars). We also saw significantly decreased false positive PSMs with increasing number of sections used. Overall, the best balance between PSM sensitivity and false positive reduction was observed when using 10 sections.

Use of human entrapment database helped in defining false-positive PSMs which were present even after utilizing the 1% global FDR cut-off. The false-positive PSMs identifications (human PSMs) from traditional large database search and traditional two-step method were evaluated and compared. Traditional large database search was able to restrict the human PSM identification rate at 1% (in line with the 1% global FDR estimation by the target-decoy method used in the PSM filtering), identifying 102 PSMs from human protein sequences. However, the traditional two-step method had a higher rate of human PSMs (2.8%), indicating an inflated false-positive rate even at an estimated global 1% FDR.

As a comparison, we performed sectioning method in three different ways, dividing the full database (*Pfu* plus human entrapment sequences) into two, five, or 10 equal-sized sections respectively. In all cases, the sectioning method yielded a more number of true positive PSMs (matches to *Pfu* sequences) when compared with the traditional large database search and controlled the number of false-positive PSMs (matches to human sequences) compared to the traditional two-step method [**Figure 3b**]. When evaluating the effect of the number of database sections used, ten sections best minimized false PSMs compared the other two (two and five sections), while maintaining a similar number of true positive PSMs as compared to the standard approach using the *Pfu* sequence database alone. These results established that sectioning of the database serves to minimize false positive PSMs, with more sections providing a more dramatic decrease, while still maintaining sensitivity for true positive PSMs – at least for a somewhat simple database containing a small number of true positive sequences (*Pfu*) compared to a large number of entrapment sequences (human).

### SIHUMI dataset

With evaluation results in hand using the single organism data from *Pfu*, we next sought to evaluate the sectioning method on a more complex dataset, one which more closely mimics the situation encountered in metaproteomics, while still offering an assessment of sensitivity and false-positive rates. Here, we used the SImplified HUman Intestinal MIcrobiota (SIHUMI) dataset^39^. The publicly available dataset was recently generated for the purpose of evaluating new methods related to metaproteomics analysis^39^. The SIHUMI dataset was generated from proteins extracted from eight microorganisms (*Anaerostipes caccae, Bacteroides thetaiotaomicron, Bifidobacterium longum, Blautia producta, Clostridium butyricum, Clostridium ramosum, Escherichia coli, Lactobacillus plantarum*) grown in a bioreactor. We first conducted a baseline standard database search against the available database^39^ composed of protein sequences from only the eight organisms contained in the sample. Results from the standard search yielded 50,651 PSMs at 1% global FDR.

We next used the traditional large database search, traditional two-step, and sectioning methods for analysis of the SIHUMI dataset. Here, we appended protein sequences from Archaea to the sequences from the eight organisms contained in the sample, with the Archaea sequences acting as a large entrapment database (see Methods). This mimics the situation in many metaproteomics analyses, where a relatively small proportion of organisms in the database are expressing proteins, while most of the sequences in the database are not contained in the sample. In this case, PSMs to the Archaea protein sequences represent false-positives, while PSMs to the eight organisms are true-positives. Results from the traditional large database search showed a loss of more than 10% true PSMs (44,375 true-positive PSMs) compared to the results from searching the database containing only the eight organisms. Meanwhile, the traditional two-step method was able to recover this number and yield 50,246 true-positive PSMs. Sectioning methods, with 5, 10, 20, and 30 sections, were also able to increase the number of true-positive PSMs (48313, 47927, 47564, and 47232, respectively) when compared with the numbers from the traditional large database search [**Figure 4a**].

**Figure 4:**
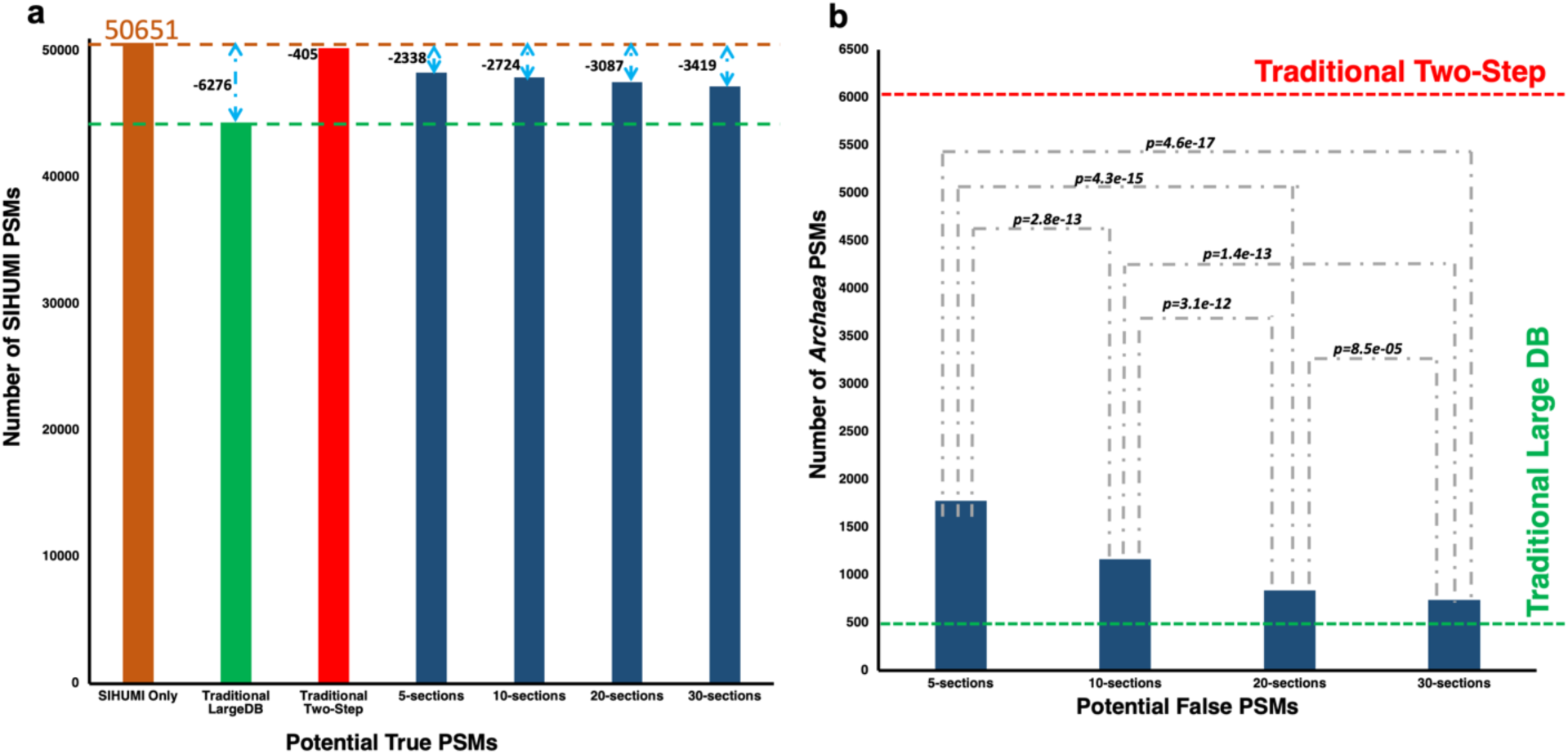
Observations using SIHUMI data, where we considered PSM(s) that matched to a SIHUMI or cRAP protein as true, while those matching to an Archaea protein were considered as false. (a) 50561 PSMs were identified when LC-MS/MS data was matched against SIHUMI + cRAP database. LC-MS/MS data when matched against an entrapment database (SIHUMI + cRAP + Archaea protein sequences) using the Traditional Large database search showed a decrease in true PSMs by more than 10%. The Traditional Two-Step method recovered almost all of these PSMs. The sectioning method also recovered a significant number of PSMs. (b) The Traditional Large database search was seen to control the false identifications (PSMs matching to Archaea protein, green dashed line), whereas the Traditional Two-Step increased the identification of false PSMs by over an order of magnitude (red dashed line). Using the sectioning method, we observed a significant decrease in the false PSM identifications when compared with the Traditional Two-Step method. This effect was greater with increasing number of sections used.

Evaluation of false-positive PSMs, those matched to the Archaea proteins, revealed that the traditional large database search method yielded about 1% false PSMs at 1% global FDR cutoff, whereas the traditional two-step method inflated the false PSMs to more than 11% [**Figure 4b**]. The sectioning method with 30 sections was again able to control the number of false-positive PSMs at around 1.5% when evaluated at 1% global FDR. We also noticed that in general the Archaea PSMs had a trend of scoring slightly worse in terms of the PSM confidence score assigned by PeptideShaker compared to most of the matches to the proteins from the eight organisms within the SIHUMI sample. As such, applying a slightly more stringent score cutoff can be used to control the global FDR and reduce the false-positive PSMs to 1%, determined by PSM matches to Archaea sequences. For example, for the 30-section results increasing stringency to a PSM score of 85 reduced the observed false-positive rate to 1% [**Figure 5**].

**Figure 5:**
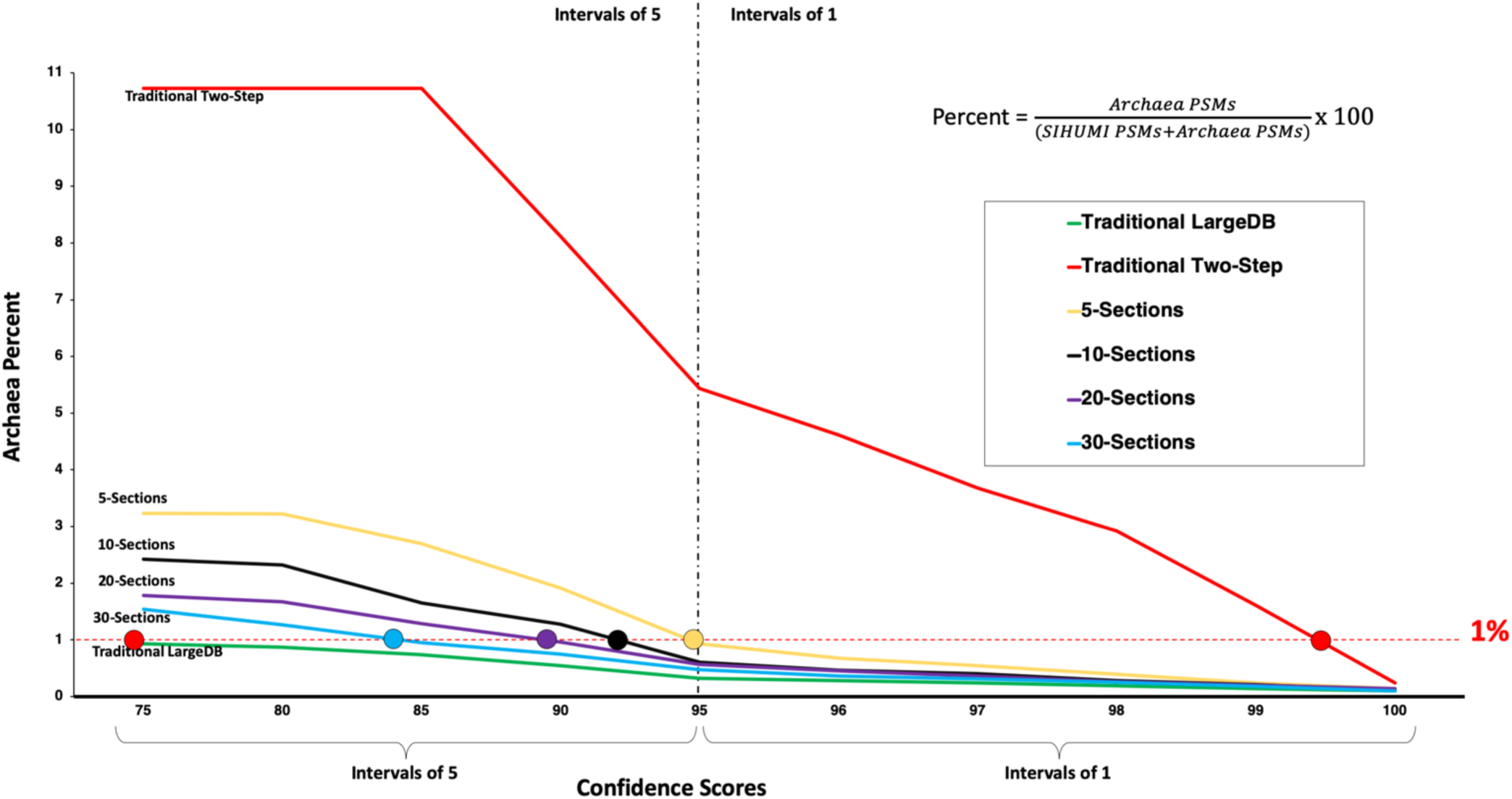
Percentage of false positive identifications in the SIHUMI dataset with corresponding PSM confidence scores. Based on an estimated global FDR of 1%, we expected matches to false sequences (derived from Archaea proteins) to approximate this rate of false positives. We observed that Traditional Large database search showed identification of about 1% false PSM identifications. The Traditional Two-Step method produced a rate 10% of false positive PSMs matching Archaea sequences, such that a very strict PeptideShaker confidence score cutoff of higher than 99 will be required to control the false positive rate. Meanwhile, the sectioning method yielded slightly higher false positives compared to the Traditional Large database search, but significantly lower compared to Traditional Two-Step method. The observed false positive PSMs also decreased as the number of sections were increased. We also observed that most of the PSMs to Archaea were assigned slightly lower confidence scores by PeptideShaker compared to those true positive PSMs. Thus, by slightly increasing stringency of qualifying PSM scores (e.g. setting a threshold of 85), the false positive rate could be significantly reduced while minimally sacrificing true positive PSMs.

**Figure 6** shows results from a deeper exploration of the relationship between global FDR, observed false positive PSMs, and sensitivity for true positive PSMs. As shown in **Figure 6**, when using a cut-off of 0.60% global FDR we were still able to identify 46,771 true PSMs while bringing the observed false-positive identifications down to 0.98% based on hits to the Archaea sequences. At 0.65% global FDR, observed false-positive identifications were 1.05%. Increasing the stringency decreased the number of true-positive PSMs only a negligible amount. Decreasing the global FDR from 1% to 0.60% global FDR, reduced the true PSMs by 1.7%, whereas it dropped the false PSMs by 38% (see Supplementary Table S1).

**Figure 6:**
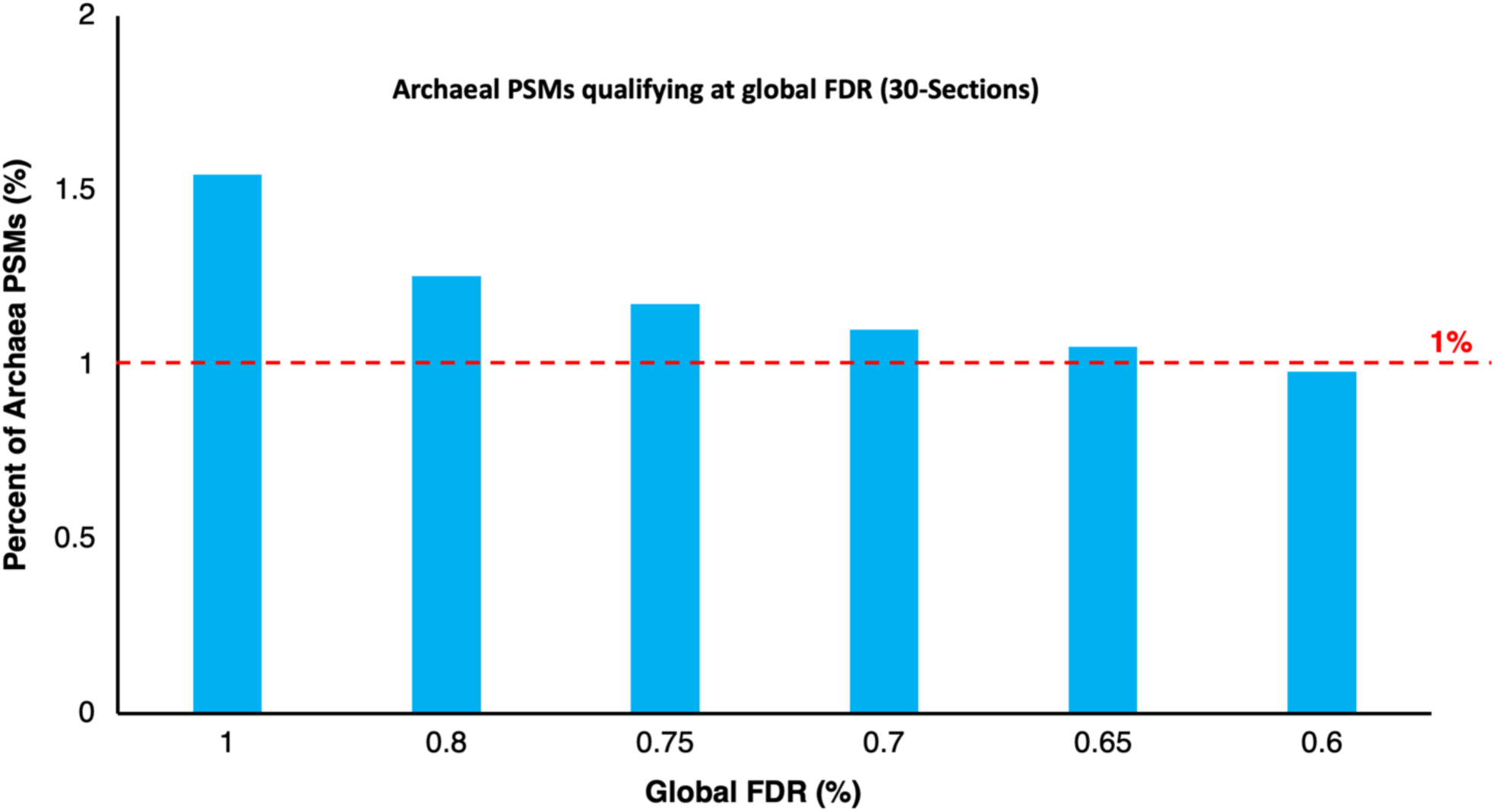
Percentage of Archaea PSM identifications at a given target-decoy global FDR for the SIHUMI dataset (30-sections). At 1% global FDR (red dashed line), the sectioning method had a 1.5% rate of PSMs to Archaea sequences. Using a slightly stricter global FDR cutoff helps in decreasing the number of false-positive identifications, for example reaching below 1% (0.98 %) false-positives at 0.60% global FDR.

Another observation related to these results was the different sizes of sequence databases uses for the generation of PSMs, depending on the method employed. The traditional large database search used a database containing 1.47 million sequences, whereas the enriched database used by traditional two-step method contained only 17,179 sequences. Meanwhile, the enriched databases generated by the sectioning method (using 5, 10, 20, or 30 sections) contained 238,191, 396,255, 578,785, and 723,243 sequences, respectively. The size of the database used inversely corresponded to the number of false-positive PSMs [**Figure 7**].

**Figure 7:**
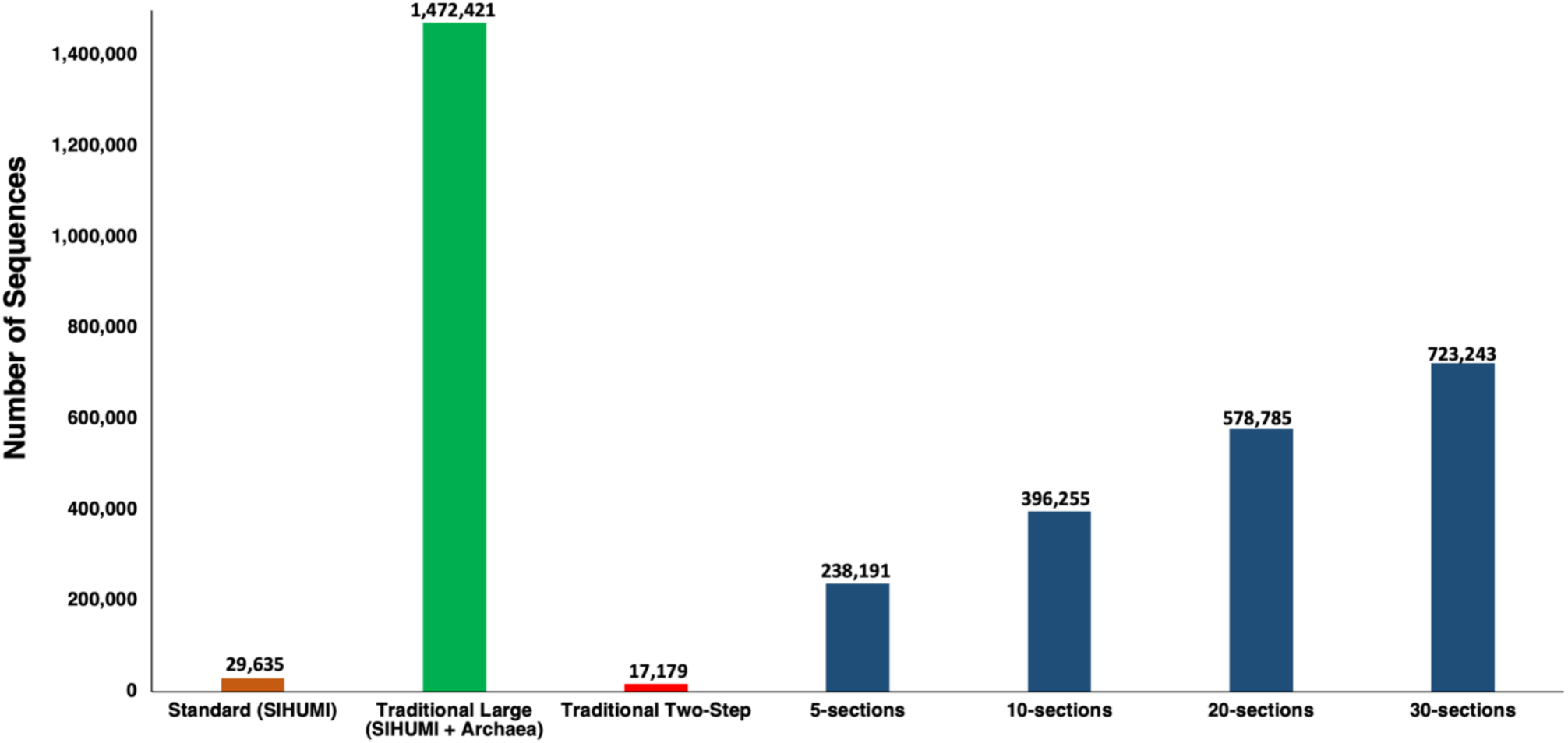
Database size used for each method for the SIHUMI dataset and its importance. The ideal protein sequence database for matching this SIHUMI dataset would include known protein sequences from the 8 organisms (29,635). To mimic a metaproteomic database, an entrapment database (1.47 Million sequences) was created by adding Archaea protein sequences. The Traditional Two-Step method reduced the database size very significantly, but to an extent such that false positive PSMs increased significantly. The sectioning method generated an enriched database of reduced size compared to the starting large database, but also contained a significant amount of proteins not detected in the sample to help control false positives. Utilizing more sections also increased the size of the database accordingly, contributing to the reduction in false-positive PSMs observed with increasing sections used.

We also tested whether or not the exact composition of the database sections used in the sectioning method had an effect on results. As described in the Methods above, this method sections the large database into randomly composed databases of a specific size. We performed ten repetitions of 5, 10, 20, and 30 sections, each time randomly assigning sequences from the large database to each section. Consistent results were observed across repeated analyses with very low coefficient of variation [**supplementary Figure S1**], demonstrating that our initial findings with the sectioning method were not due to an artifact related to database composition.

### Glacial meltwater dataset

After the evaluation of the sectioning method using model datasets, we next used it on a complex microbiome dataset, representative of the scope of most MS-based metaproteomics studies. This dataset was derived from a protein sample extracted from a microorganism community contained in the meltwater from a glacier^40, 41^. The protein database was generated using SixGill^20^ and MOCAT^22^ tools using the whole genome sequencing of the metagenomes extracted from the same meltwater samples. The database contained over five million protein sequences. We compared the results from the traditional large database search (matching the complete 5 million sequences database to MS/MS data), with the results from the sectioning method, using 5-sections. The sectioning method helped to reduce the database size to 204,608 sequences in the enriched database.

We were able to identify 20% more microbial PSMs from the meltwater sample (excluding common contaminants), at a global FDR of 1% when compared with the similar output from the traditional large database search. We used all the peptides from these PSMs to perform Unipept 4.0 analysis^42^, which helps in assigning metapeptides to specific taxonomic groups, functional classes, and protein-level enzyme commission (EC) numbers. The increased PSMs from the sectioning method resulted in increased assignment across all categories (taxonomy, functional classes, EC numbers) compared to the traditional method. The results from the sectioning method was able to identify most of the terms identified by the traditional method, showing substantial overlap [**Figure 8**]. Analysis of the assigned taxa, functions and EC numbers that were found by both methods revealed that sectioning increased the depth of data supporting these assignments [**Figure 8**]. Put another way, the increased the number of PSMs from sectioning provided more confidence in taxa and functional group assignments reported by Unipept, providing deeper insight into interpretations of the results.

**Figure 8:**
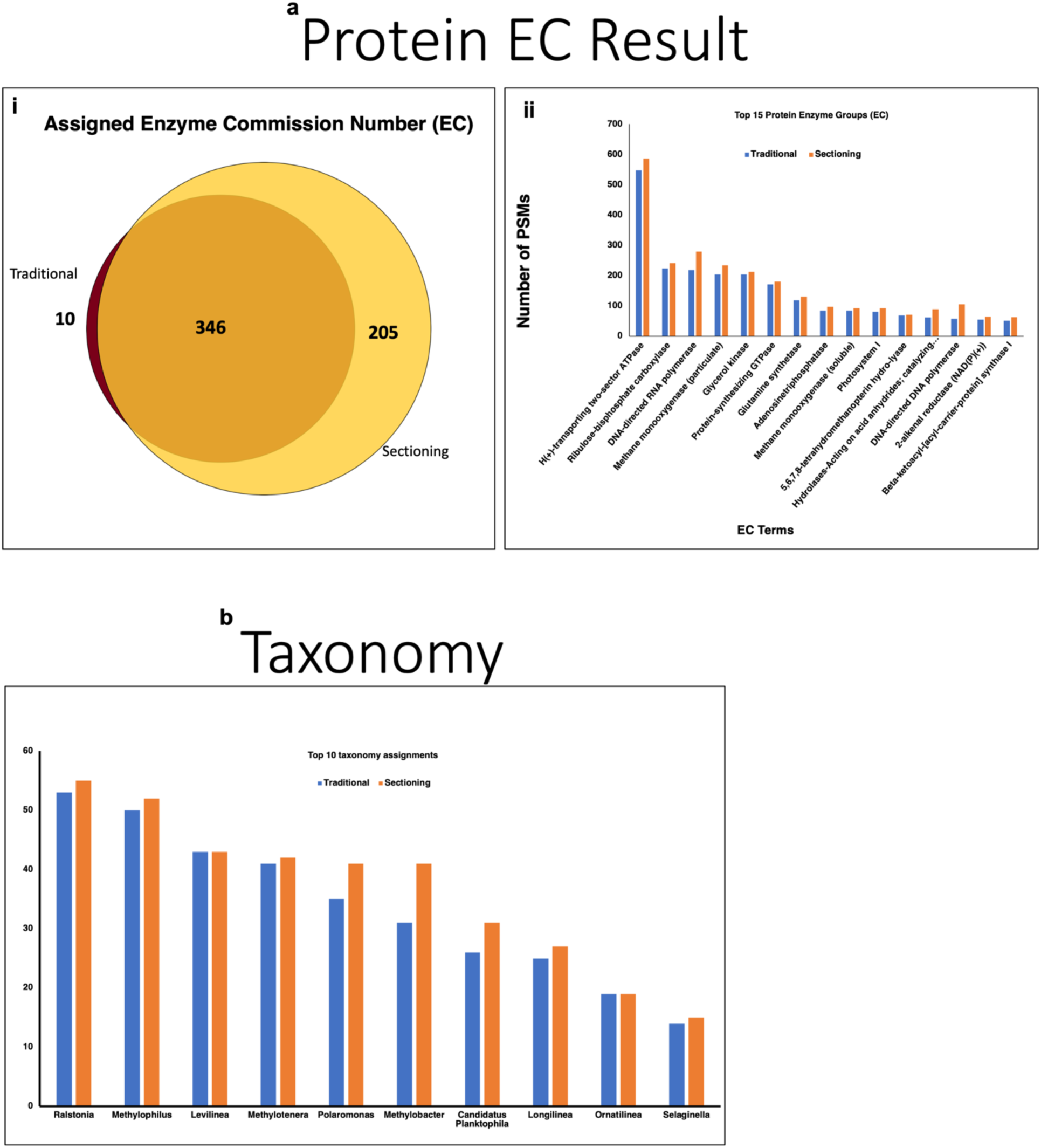
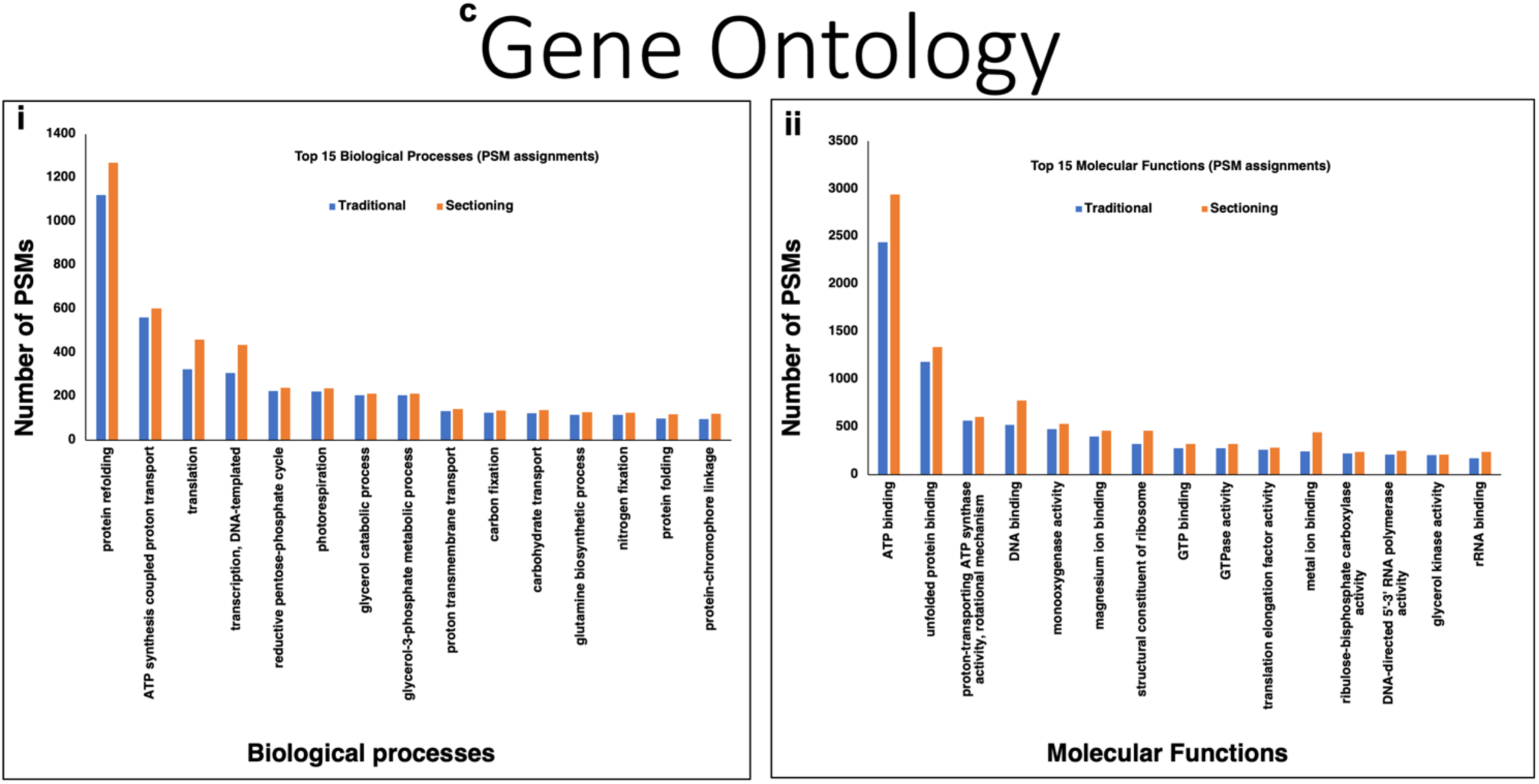
Meltwater dataset: Comparison of Unipept outputs from sectioning method and traditional large database searching method. (a) Protein enzyme commission (EC) assignment report from Unipept shows that the sectioning method was able to assign majority of the EC numbers identified by Traditional Large database searching method, as well as some additional EC numbers. For the sectioning method, more PSMs were assigned to each EC number compared to the Traditional Large database searching method, improving the confidence in the assignments. (b) The sectioning method was able to identify more taxonomy groups using Unipept compared to the Traditional Large database searching method. Also, more PSMs were assigned to each of the top ranking taxonomy groups when sectioning method was used, improving confidence. (c) Similar to EC number and taxonomy, PSMs from the sectioning method were assigned to more gene ontology groups, both biological processes (i) and molecular functions (ii). Also, more PSMs were assigned to each of the top-ranking gene ontology terms in both biological processes and molecular functions when the sectioning method was used.

## Discussion

To evaluate our sectioning method, we used model MS/MS datasets (*Pfu* and SIHUMI). These datasets mimicked the situation encountered in larger databases used in metaproteomics applications, where a relatively small proportion of protein sequences within the database are detected by the mass spectrometer and contained within the collected MS/MS data. We first matched the MS/MS spectra to the standard organism-specific protein databases and then compared these results to these same databases appended to a much larger entrapment database. As others have shown^33, 34^, when including a large entrapment database the sensitivity for detecting PSMs to the known proteins within the samples decreases when using traditional sequence database searching methods [**Figure 3, 4**]. This sensitivity loss is a concern for metaproteomics, or any application where large protein sequence databases are employed (e.g. generating protein databases for proteogenomics).

We then tested the traditional two-step method^33^ that generated an enriched, much smaller database for matching MS/MS data. We observed that in both the datasets, the traditional two-step method helped in increasing the sensitivity, generating an increased number of PSMs passing the 1% global FDR threshold. However, our tests also confirmed that the enriched database generated by the traditional two-step method overcompensates, producing a database so small that it increases the potential for false positive PSMs significantly (even at 1% global FDR estimated using standard target-decoy methods) [**Figure 5, 7**]. Our prior publications on the two-step method recognized this potential for increased false positives and suggested utilizing post-processing filtering and spectral quality validation of PSMs to further ensure accuracy^33, 43^. Here, we have confirmed the false-positives are a real concern with the traditional two-step method.

We developed the sectioning method to address the limitations above -- the sensitivity loss when using a traditional database searching methods against a large database, and the observed increase in false-positive PSM when using the traditional two-step method. Therefore, the sectioning method is an improvisation of the traditional two-step method, with an intention to find a method that balances sensitivity while controlling for false-positive PSM identifications.

The results from the Pfu and SIHUMI datasets both showed the benefit of our sectioning method. For both datasets, generating an enriched database from sequence database searching of sections increased the sensitivity compared to the traditional large-database method, while significantly decreasing false-positives compared to the traditional two-step method. The magnitude of this effect was a bit less for the Pfu dataset (derived from a single organism with a simple proteome) compared to the more complex SIHUMI dataset (derived from eight model organisms), suggesting that the benefit of the sectioning method may increase with size and the complexity of the database being used. It was also clear that dividing the initial database into more sections helped control false-positive PSMs more effectively [**Figure 9**]. Not coincidentally, the enriched database size also increased with more sections used, likely contributing to the reduced false-positive rate by including more competing protein sequences for MS/MS matching, leading to better scoring separation between false PSMs and true PSMs. However, the enriched database is still significantly smaller than the initial large database, such that PSM sensitivity is also maintained. Therefore, our results indicate the sectioning method offers an optimal balance in database size to provide sensitive PSM identification while controlling false-positives.

**Figure 9:**
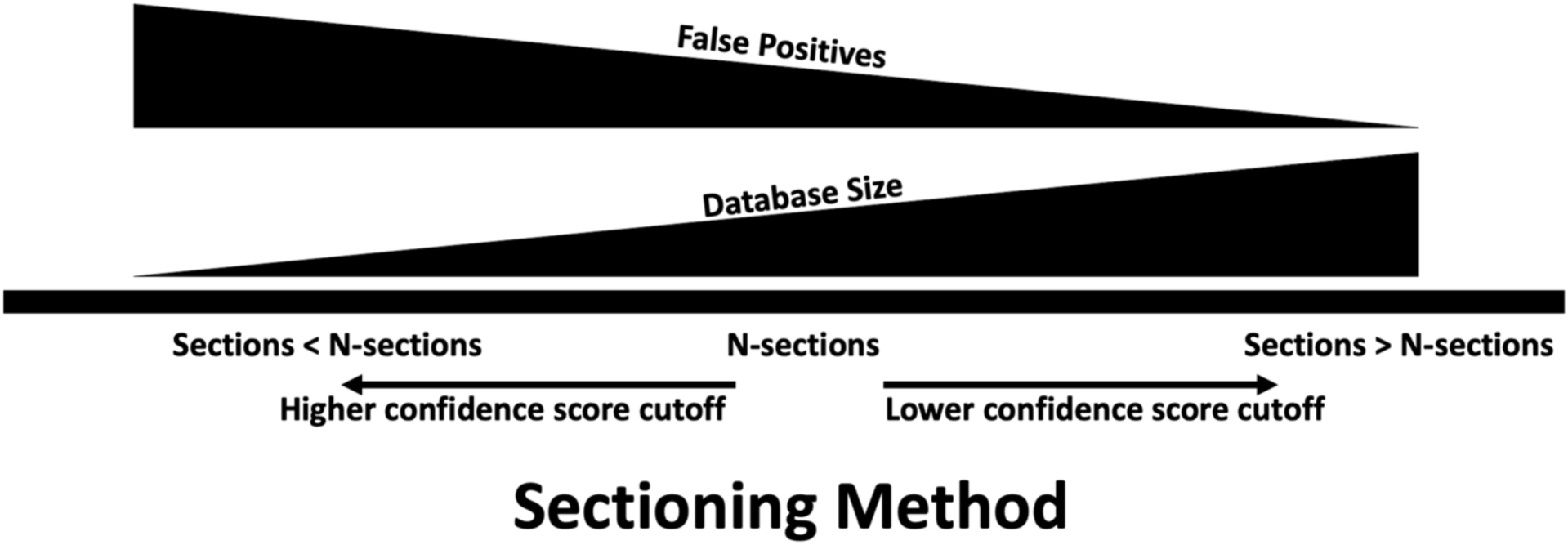
A summary of the findings from the evaluation of the sectioning method for metaproteomics databases. While increasing the number of sections decreases the false-positive potential, it comes with the trade-off of generating enriched databases of increasing size. PSM scores of lower quality but passing the 1% global FDR threshold can be assumed as true with more sections. Meanwhile, a lower number of sections provides a smaller-sized enriched database, but with a cost of increasing potential for false positive identifications. Stricter cut-offs of PSM scores and/or stricter global FDR thresholds (< 1%) is suggested.

**Figure 7** shows this effect of number of sections on the enriched database size. The more sections used means a smaller number of sequences in each section and a more restricted search space for the first round of matching MS/MS to these sequences. Using our permissive scoring cutoffs in the first round of matching of MS/MS spectra to each section leads to more proteins qualifying as potential members of the samples when using more sections. As such, more sections lead to an overall larger enriched database for the second step of sequence database searching. However, as shown in **Figure 7**, even the larger enriched databases generated with more sections are still significantly smaller than the initial database.

Despite the demonstrated ability of the sectioning method to significantly decrease the observed false-positive PSMs compared to the traditional two-step method, we did observe a slightly increased potential for false-positives. For example, even with the best-case scenario of 30-sections, the SIHUMI dataset showed a measured false-positive rate of about 1.5% when counting the PSMs to Archaeal peptides, which is higher than the estimated global false positive of 1% using the target-decoy method [**Figure 5**]. When analyzing these PSMs to Archaea more closely, we did notice the trend that the false-positive PSMs generally had lower scores as assigned by PeptideShaker, indicating that these are slightly lower confident data, even though they passed the 1% global FDR threshold. As such, when employing the sectioning method, we would suggest that extra stringency be considered for qualifying PSMs, such as using a lower global FDR to effectively increase stringency on PSMs qualifying as correct. As we showed in **Figure 6** and also Supplemental Table S1, lowering the global FDR threshold to a value such as 0.6% reduced the observed false-positive PSMs to below 1% while only minimally decreasing the sensitivity of detecting true-positive PSMs.

Some recent studies have focused on quantification of metaproteomics datasets^44^. However, software suites that can perform quantification along with identification (such as MaxQuant^45^ MaxLFQ^46^), function optimally only when intermediate or small database size is used. In such cases, the number of sections used could be selected in order to maintain an enriched database of manageable size, with the caveat that there may be an increased potential for false-positives with a smaller database. This might require an additional step of filtering peptides/PSMs at a lower global FDR as described earlier. However, for quantification methods that are independent of identification (such as moFF^47^ and FlashLFQ^48^), and do not have constraints on database size, a larger number of sections would be recommended to control for false positives.

After evaluating and understanding the benefits of the sectioning method on model datasets (*Pfu* and SIHIMU), we applied the method to data from a representative metaproteomics study of microbial proteins expressed in a glacier meltwater sample. Here, we observed a significant decrease in the size of the protein sequence database used for MS/MS matching when using the sectioning method to generate the enriched database (reduction from five million sequences to about 200,000 sequences). Accordingly, this enriched database enabled a significant increase in the identified PSMs when compared with the results from the traditional database search. We were encouraged that a vast majority of the additional PSMs identified using the sectioning method mapped to the same taxa and functional classes (determined by Unipept analysis) as those initially identified using the traditional database searching method. This supports the assertion that the sectioning method is adding accurate PSMs, which provide deeper coverage and more confident assignments of taxa and functional pathways represented by the metaproteomics data. As such, we are confident that the sectioning method will provide improved information from diverse applications of metaproteomics.

Proteogenomics, where customized protein sequence databases are generated from genomic or transcriptomic data, also suffers from issues related to large databases for MS/MS matching^5, 49^. The sectioning method described here should also benefit proteogenomic applications. However, proteogenomics has some unique features different from metaproteomics, such as the need to match MS/MS to specific classes of sequence variants and estimate class-dependent FDR values. Therefore, demonstrating the value of sectioning for proteogenomics will take a dedicated evaluation and optimization study of its own.

Finally, implementing the sectioning method requires multiple complex steps, including *“n+1”* number of database searches when using *“n”* sections, as well as the need to aggregate results to generate the enriched database for further matching to MS/MS. In order to facilitate the use of this method and make this method accessible to the community, a workflow automating these steps, with default parameters defined, in a usable platform is a necessity. As such, we have developed this method in the flexible Galaxy platform^50^, as part of the Galaxy for proteomics (Galaxy-P) development project. The complete and automated workflow can be accessed from the European Galaxy public server (https://usegalaxy.eu/u/galaxyp/w/sectioningworkflowgalaxyp) and Metaproteomics gateway^51–53^ (http://129.114.16.192/u/pravs/w/sectioningworkflowgalaxyp). Supplemental Information provides instructions on accessing and using this workflow, along with demonstration input data.

## Conclusion

We have demonstrated the value of using database sectioning for large database searching, with a focus on metaproteomic applications. Coupled with the accessibility to the automated workflow and tools necessary to carry-out this method within Galaxy, we believe that this method should provide researchers a highly valuable approach to generate improved results in large database applications in MS-based proteomics – from metaproteomics and beyond.

## Methods

### Sectioning Method overview

Database sectioning method was implemented as a workflow on Galaxy platform. It can, however, be implemented on other platforms as well. Based on the complexity and number of steps and tools required, it was an ideal fit for implement within Galaxy, where it can be automated and made accessible to others. The Sectioning method, as illustrated in [**Figure 1**], divides large databases into more manageable sections, each of which are matched to the MS/MS data. Any sequences matched, even at very low score and without applying any FDR filtering step, are mapped to their inferred proteins and used to generate a reduced database that is enriched for proteins most likely present in the sample. In addition to the selected sequences, equal number of randomly selected sequences from the original database are also added to the enriched database. The randomly selected sequences help to provide sufficient background “noise” proteins that are not detectable in the sample and allow for improved FDR evaluation. For example, if “*x*” number of sequences were selected through first-step search, “*x*” number of additional randomly selected protein sequences are selected from the original large database and included in the enriched database. The enriched database is then matched to the MS/MS dataset for generating PSMs. The sectioning method workflow can be accessed through European Galaxy Server (https://usegalaxy.eu/u/galaxyp/w/sectioningworkflowgalaxyp), Metaproteomics Gateway^51–53^ (http://129.114.16.192/u/pravs/w/sectioningworkflowgalaxyp). Workflow shared here can be downloaded as GA file (.ga) and can be uploaded onto any other Galaxy instance provided all the required software tools are installed on that instance. Also provided in the Supplemental Information is the instructions on accessing this workflow. For sequence database searching and PSM generation and scoring, this workflow uses SearchGUI^54^ and PeptideShaker^55^. These two programs were used for the evaluation of the sectioning method as described below.

For *Pfu* dataset, we used SearchGUI^54^ (SG) (version 3.2.13) and PeptideShaker^55^ (PS) (version 1.16.9) to match the MS/MS spectra with FASTA databases along with contaminants from cRAP database^56^. Although SG has the option to use as many as 9 search algorithms, but for this evaluation purpose, only four search algorithms (X!tandem, OMSSA, MSGF+, and Comet) were used.

Search parameters for the *Pfu* dataset used were trypsin digestion where two missed cleavages were allowed. Carbamidomethylation of Cysteine was selected as a fixed modification. Oxidation of methionine, phosphorylation of serine, threonine, and tyrosine were selected as variable modifications. The accepted precursor mass tolerance was set to 10 ppm and the fragment mass tolerance to 0.5 Da with minimum charge as 2 and maximum charge as 4. Peptide length ranging from 8 – 50 amino acids were used for filtering in PeptideShaker.

For SIHUMI dataset, we used SearchGUI^54^ (SG) (version 3.3.10) and PeptideShaker^55^ (PS) (version 1.16.36.2). Four search algorithms (X!tandem, OMSSA, MSGF+, and Comet) were in SG. Search parameters for the SIHUMI dataset used were trypsin digestion with two missed cleavages was allowed. Carbamidomethylation of Cysteine was selected as a fixed modification and methionine oxidation was selected as a variable modification. The accepted precursor mass tolerance was set to 5 ppm and the fragment mass tolerance to 0.02 Da with minimum charge as 2 and maximum charge as 4. For Peptide Shaker, the accepted peptide length ranged from 8 - 50 amino acids.

For meltwater dataset, we used SearchGUI^54^ (SG) (version 3.3.10) and PeptideShaker^55^ (PS) (version 1.16.36.2). Four search algorithms (X!tandem, OMSSA, MSGF+, and Comet) were in SG. The search parameters for the meltwater dataset were trypsin digestion with three missed cleavages were allowed. Carbamidomethylation of Cysteine was selected as a fixed modification. Oxidation of methionine and deamidation of glutamine and asparagine were selected as a variable modification. The accepted precursor mass tolerance was set to 10 ppm and the fragment mass tolerance to 0.02 Da with minimum charge as 2 and maximum charge as 6. For Peptide Shaker, the peptide length ranging from 8 - 50 amino acids were accepted.

### Sectioning Method Evaluation

For evaluation of the sectioning method, we used two standard datasets (*Pyrococcus furiosus* and SImplified HUman Intestinal MIcrobiota datasets). Both provided “ground truth” data for matching PSMs to organism-specific proteins. In order to accurately assess the identification of false PSMs, entrapment databases were created for both the datasets (see methods below). As a first baseline standard step, the MS/MS datasets were matched with the corresponding standard protein sequence database that contained proteome sequences of the expected organism(s) and contaminants. The numbers from this standard search was further used to compare it with the results from the other methods, traditional large database search method, traditional two-step method, and sectioning methods. Target-decoy method was used to evaluate global FDR cutoff and to qualify PSMs for comparison. PSMs qualifying at a global FDR cutoff of 1% and resulting from the expected organism’s proteome were considered as true positives, whereas those PSMs qualifying at the global 1% FDR cutoff matching to proteins in the entrapment proteome were considered false positive identifications.

### *Pyrococcus furiosus* (*Pfu*) data and creation of entrapment database

Pfu dataset was used from a previously published dataset^37^, downloaded from the publicly shared files on EBI-PRIDE Archive (Project ID: PXD001077). Proteome sequences of *Pfu* was downloaded from Uniprot and contained 2045 unique protein sequences). Contaminants (116 sequences) [Ref: ^56^] was added to the *Pfu* proteome sequences using Galaxy tool called “Protein Database Downloader” (https://github.com/galaxyproject/tools-iuc/tree/master/tools/dbbuilder).

The entrapment database was created by adding human proteome sequences (93,502 sequences) to the *Pfu* sequence and contaminant sequences. Human proteome sequences were downloaded from Uniprot database using Galaxy tool “Protein Database Downloader”.

### SImplified HUman Intestinal MIcrobiota (SIHUMI) data and creation of entrapment database

Proteins from eight microorganisms (*Anaerostipes caccae, Bacteroides thetaiotaomicron, Bifidobacterium longum, Blautia producta, Clostridium butyricum, Clostridium ramosum, Escherichia coli, Lactobacillus plantarum*) grown in a bioreactor were extracted to generated SIHUMI dataset. The raw file can be accessed from (https://files.ufz.de/~molsyb-ims-2018/ORNL_Easy_nLC1200_contest2_C1_2ug_UHPLC_gradient240min_ac50cm.raw) The dataset was publicly shared at 3rd International Metaproteome Symposium (https://www.ufz.de/index.php?en=44639). The proteome sequences of these eight microorganisms were also shared through on the symposium page (https://www.ufz.de/export/data/2/211671_Contest_Sample_1_SIHUMI.fasta). The protein sequence database was then merged with contaminant sequences and the archaeal proteome sequences (containing 1.44 million sequences downloaded from the NCBI-NR database). This merged database was used as an entrapment database where any PSM matching SIHUMI or contaminants were considered as true positive identifications whereas those matching to Archaea sequences were considered false positives.

### Metaproteomic meltwater data

The protein samples were extracted from the microbiome of meltwaters on the surface of and underneath a glacier^40, 41^. For our work, data from a single collection timepoint was used. The collection of metaproteomic MS/MS data files (accessible from https://arcticdata.io/catalog/view/doi:10.18739/A2VX06340) were used to match against a protein sequence database generated from whole genome metagenomics data derived from the same meltwater sample. Metagenomes were assembled using MOCAT^21^ and then translated into protein sequences. Additionally, the SixGill tool^20^ was used to generate an alignment-free metapeptide sequence database. A merged protein sequence database was used containing protein sequences from MOCAT and SixGill, as well as common contaminants detected in MS-based proteomics experiments. The protein sequence database contained 5,117,895 sequences. MS/MS data was first matched with complete 5.1 million sequence (traditional method) and then sectioning method was used – using 5 equal sized sections of approximately 1.02 million sequences each – to create an enriched database. All PSMs to peptides, excluding contaminant peptides, from both traditional and sectioning method were further analyzed through Unipept^42^ (https://unipept.ugent.be/datasets). Unipept allowed assignment of identified peptides to enzyme commission numbers (EC numbers), taxonomy, and gene ontology terms like biological processes and molecular functions.

## Acknowledgements

We would like to thank European Galaxy and Freiburg Galaxy team for providing the computational resource and storage for the Galaxy implementation and sharing of the database sectioning method. We would also like to thank the Minnesota Supercomputing Institute (MSI) and Jetstream for computational resources.

We acknowledge funding for this work from the grant National Cancer Institute - Informatics Technology for Cancer Research (NCI-ITCR) grant “1U24CA199347”, National Science Foundation (U.S.) grant “1458524”, and a grant through the Norwegian Centennial Chair (NOCC) program at the University of Minnesota to T.G.

We would also like to acknowledge the Extreme Science and Engineering Discovery Environment (XSEDE) research allocation BIO170096 to P.D.J. and use of the Jetstream cloud-based computing resource for scientific computing (https://jetstream-cloud.org/) maintained at Indiana University. We also acknowledge the support from the Minnesota Supercomputing Institute for maintenance and update of the Galaxy instances. The European Galaxy server that was used for some calculations is in part funded by Collaborative Research Centre 992 Medical Epigenetics (DFG grant SFB 992/1 2012) and German Federal Ministry of Education and Research (BMBF grants 031 A538A/A538C RBC, 031L0101B/031L0101C de.NBI-epi, 031L0106 de.STAIR (de.NBI)).

