## Supplementary Information for "A sectioning and database enrichment approach for improved peptide spectrum matching in large, genome-guided protein sequence databases"

### Supplementary/Supporting Information

#### Supplementary Section 1: Supplementary Figure

Figure S1

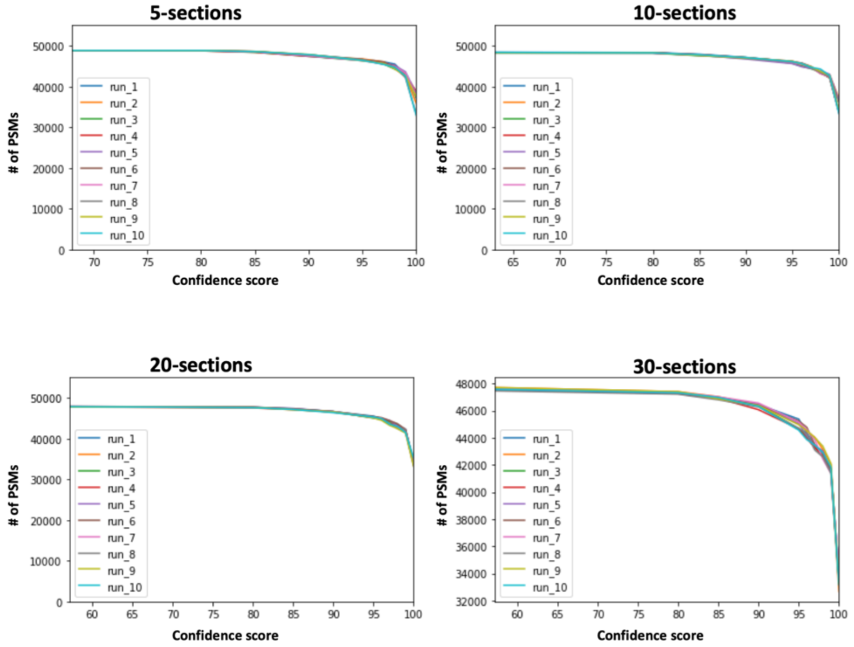


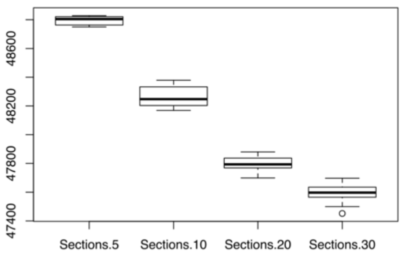


| | Coefficient of variation(CV) |
|---|---|
| 5-sections | 0.00062 |
| 10-sections | 0.00153 |
| 20-sections | 0.00120 |
| 30-sections | 0.00161 |

Figure S1: SIHUMI PSMs: Sectioning involves randomly assigning protein sequences in each sectioned database bin. Shown here are results from running 10 such randomizations for each 5, 10, 20, and 30 sections. Figures indicate that each randomization yielded very similar results, indicated by a low coefficient of variation shown in the table.

**Supplementary Section 2: Supplementary Table**

Table S1

| Global FDR cutoff | False PSMs | True PSMs | False positive PSM (%) | (%) Decrease in False PSMs, relative to the 1% global FDR level | (%) Decrease in True PSMs, relative to the 1% global FDR level |
|---|---|---|---|---|---|
| **1** | 747 | 47581 | 1.55 | NA | NA |
| **0.8** | 600 | 47245 | 1.25 | 19.68 | 0.71 |
| **0.75** | 561 | 47176 | 1.18 | 24.90 | 0.85 |
| **0.7** | 524 | 47065 | 1.10 | 29.85 | 1.08 |
| **0.65** | 499 | 46943 | 1.05 | 33.20 | 1.34 |
| **0.6** | 463 | 46771 | 0.98 | 38.02 | 1.70 |

Table S1: Decrease of True and False PSM identifications at a given target-decoy global FDR for the SIHUMI dataset (30-sections).

#### Supplementary Section 3: Accessing Sectioning Workflow

The workflow can be accessed from the Galaxy server hosted publicly from Europe (https://usegalaxy.eu/u/galaxyp/w/sectioningworkflowgalaxyp).

To save the workflow, click on the "+" sign found on the right top corner of the page.

The workflow requires two inputs – dataset collection of MGF files and protein sequences in FASTA format.

Core parameters and test dataset

1) Tool "Split file": set "Number of new files" to number of sections needed for analysis.
2) SearchGUI and PeptideShaker parameters should be checked and modified based on the dataset and its acquisition method. SearchGUI and PeptideShaker are used two times in the workflow. Modify parameters at both times as required.
3) Query Tabular (the one reading "Extended PSM Report" from PeptideShaker): The SQLite query can be modified depending on the optional score cutoff required.
   "select distinct Proteins from psm where Proteins not like '%_REVERSED%' and Confidence >= 0"
   The "Confidence >= 0" is the most lenient and recommended cutoff. This would include Proteins identified from all PSMs. The Confidence value ranges from 0 to 100 and any value within this limit can be used. Though we recommend using lower value here.
4) A sample data is shared for testing the database sectioning method and can be accessed from a shared history on European Galaxy (https://usegalaxy.eu/u/galaxyp/h/sectioningsampleinputdata).
